# Multiple Confounders Correction with Regularized Linear Mixed Effect Models, with Application in Biological Processes

**DOI:** 10.1101/089052

**Authors:** Haohan Wang, Jingkang Yang

## Abstract

In this paper, we inspect the performance of regularized linear mixed effect models, as an extension of linear mixed effect model, when multiple confounding factors coexist. We first review its parameter estimation algorithms before we introduce three different methods for multiple confounding factors correction, namely concatenation, sequence, and interpolation. Then we investigate the performance on variable selection task and predictive task on three different data sets, synthetic data set, semi-empirical synthetic data set based on genome sequences and brain wave data set connecting to confused mental states. Our results suggest that sequence multiple confounding factors corrections behave the best when different confounders contribute equally to response variables. On the other hand, when various confounders affect the response variable unevenly, results mainly rely on the degree of how the major confounder is corrected.

## I. INTRODUCTION

With the necessity of introducing large data sets, researchers are competing to generate enormous data sets to study. Beyond the possibility of collecting relevant information simultaneously, batches of data have to be collected first and then integrated into one data set. This data acquisition conduct does not raise obvious hazards for traditional tasks such as face recognition or optical character recognition [14], [30], however, may dramatically affect the generalizability of trained models or scientific discovery followed-up on many modern tasks where data is sensitive to confounding factors [17].

Confounding factors have been discussed in statistics for a long time [2], [22], so we only briefly recapitulate the ideas here. For a study of the relationship between explanatory variable *X* and response variable *Y*, confounding effects occur when there is a third variable *Z* that influences both *X* and *Y*. Therefore, frequently, the correlations between *X* and *Y* is not intrinsic. In other words, the relationship between *X* and *Y* arises from confounding factor *Z*, rather than real associations. However, to be a factor that have real impact on the response variable, it can neither be the intermediate variable nor the one only relevant variable in a small batch of data. Thus, the discovered relationship is a spurious relationship between *X* and *Y*, which happens more than often in real-world. For example, a biological process may be correlated with the temperature or humidity of when experiments are conducted [19], or a psychological reaction may be associated with cultural or demographic information of where data is collected [31]. Similar circumstances also apply to GWAS [21], where samples in the same batch presumably share some genotypic patterns or phenotype properties, resulting in the discovered association to be a spurious one.

To effectively correct confounding effects, linear mixed model (LMM) is introduced [20]. Various works have been introduced to apply LMM to different domains. For example, Cnaan et al. focused on unbalanced repeated measures data and longitudinal data [5]. Krueger and Tian compared linear mixed model and ANOVA to derive two advantages of linear mixed model: the ability to accommodate missing data points often encountered in longitudinal datasets and the capacity to model nonlinear, individual characteristics [12]. On the frontier of bioinformatics research. Schelldorfer et al. proved that linear mixed model with Lasso is beneficial whenever a grouping structure among high-dimensional observations exist [26]. In addition, Ghidey et al. showed that the L_2_ penalization can be utilized to smooth random effects distribution [8]. Different penalizations can guide linear mixed models to exhibit different performances [23].

Although there are some previous researchers investigating the case when two or more confounders coexist in one data set, for example, LaGasse et al. worked on multiple confounding factors corresponding to fetal exposure to understand how drugs influence on the fetus [13], and Griffiths took multiple confounding effects in multiple linear regression to evaluate the effect of bulk tumor resection on survival [27], few of them inspected how linear mixed model perform with multiple confounders. Therefore, a valuable question remains unanswered about how to correct the multiple confounding factors with linear mixed model effectively.

In this paper, we mainly focus on the problem when multiple confounding factors coexist. We introduce three methods that can work with multiple confounding factors in different perspectives, ahead of which, we first review the traditional linear mixed model and present the regularized versions derived from original ones, in addition to our discussion of two different parameter learning approaches for LMM, namely *MLE* and *REML*, which could be straightforwardly generalized for regularized linear mixed models.

In our experiments, we compare the performances of traditional and regularized linear mixed models under different parameter learning likelihood functions, as well as with different multiple confounding factor correction methods. We evaluate the performance on variable selection task as well as prediction task on three different data sets: 1) synthetic data set, 2) semi-empirical synthetic data set based on real-world genome sequences, 3) real-world brain wave data set relating confusion mental state. Our results indicate that L_1_ regularized model with *sequence* multiple confounding factor correction method achieves the best result consistently.

This paper is structured as the following: we first briefly review standard linear mixed models and its parameter estimation likelihood functions, then present regularized linear mixed models with L_1_ and L_2_ regularizers. Finally, methods focusing on multiple confounding factors are introduced, followed by experiments on synthetic data and real-world data. Then we conclude this paper with a brief discussion followed by summaries and future directions.

## II. REGULARIZED LINEAR MIXED MODEL

In this section, we will introduce regularized linear mixed model, ahead of which, we first briefly discuss original linear mixed model and its parameter learning likelihood functions as background.

### A. Linear Mixed Model

Linear Mixed Model (LMM) is a model that has been widely appreciated in the effectiveness of correcting confounding factors introduced by batch effects of data collected and summarized in groups [20]. As an extension of the linear regression model, it describes the relationship between a response variable and explanatory variables, with coefficients that can vary on one or more grouping variables to correct batch effects. Correspondingly, a mixed-effects model consists of two parts, fixed effects as conventional linear regression and random effects that are associated with different batches.

Formally, for a standard linear regression, suppose we have *m* samples, with response variable *y* = (*y*_1_*, y*_2_,...*y_m_*) and known explanatory variables *X* = (*x*_1_*, x*_2_*,...x_m_*). For each *i* = 1, 2*,..., m*, we have *x_i_* = (*x_i,_*_1_*, x_i,_*_2_*,...x_i,p_*), i.e., *X* is of the size *m×p*. A standard linear regression is in the formality of *y* = *Xβ* where *β* stands for parameter vector for fixed effects.

Considering batch effects, we have linear mixed model as following:

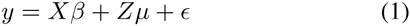

where *μ* stands for vector of random effects, *Z* stands for a designed matrix for random effects of the size *m × t* and stands for observation noise.

Equation 1 can be formalized as the following Gaussian process

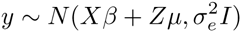

where 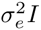 accounts for the observation noise ∈ in previous equation. Further, the distribution of random effect *μ* is assumed to be 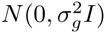 and above Gaussian process can be written as:

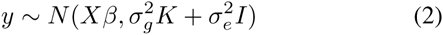

which is a standard form of LMM, where *K* = *ZZ^T^*.

#### a) Parameter Estimation

There are numerous algorithms accounting for parameter estimation of LMM. For major focus, here we focus on one state-of-art algorithm for (restricted) maximum likelihood estimation. Readers can refer to other literature for a thorough introduction of algorithms [4], [10], [20].

To begin with, from Equation 2, we could derive LMM’s log-likelihood function as following:

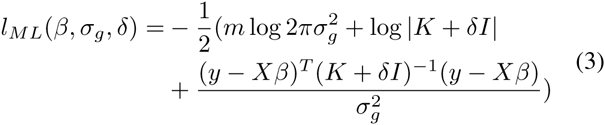

where 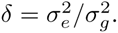

However, Equation 2 does not take into account the loss in degrees of freedom resulting from estimating fixed effects. A straight forward extension of it is Restricted Maximum Log-Likelihood, as following:

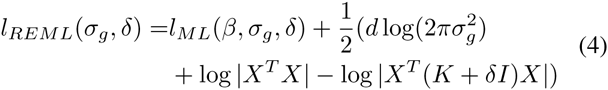

where *d* stands for the loss of degrees.

Solving these non-convex optimization problems of maximizing *l_ML_*(*β, σ_g_, δ*) with standard methods like grid search will resolve the parameter estimation problem.

A more effective way to solve Equation 3 and Equation 4 is introduced by [16], named FaSTLMM, which stands for factored spectrally transformed linear mixed model. It solves Equation 3 by taking spectral decomposition of *K* (i.e. *K* = *USU^T^*) and rewrite Equation 3 into:

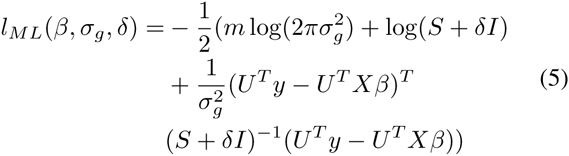

Then, *β* and 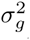 can be solved with closed form by taking derivative and setting the derivatives to zero respectively. After plugging in the closed form of *β* and 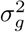 back into Equation 5,*δ* can be solved with Brent search [3].

Specifically, fixed effect sizes *β* can be achieved with:

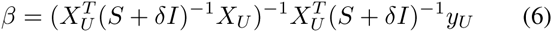

where *X_U_* = *U^T^ X* and *y_U_* = *U^T^ y*. Equation 6 offers the closed form of fixed effect sizes encoding how explanatory variables contribute to response variables.

### B. Regularized Linear Mixed Model

With the background built up, we now proceed to regularized linear mixed models. Following the same way of how regularized linear regression is extended from vanilla linear regression, we introduce two forms of regularized linear mixed models here and discuss the potential for LMM to be extended with other regularizers.

#### 1) Sparse Linear Mixed Model

One distinct advantage of Sparse Linear Mixed Model is that it results in a sparse fixed effect vectors that are corresponding to the most relevant explanatory variables associated with response variables. It is attained with sparse regularizer induced to linear mixed model during parameter learning process. One straightforward way is to replace Equation 6 as following:

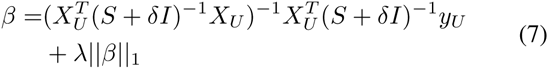

where *λ* is a prior to control the strength of sparsity regularizer. As we can see, this is a direct extension of traditional Lasso [28], also discussed in [7], [25].

Another direction to introduce sparsity into LMM, which is more intuitive, but less employed in reality, is to induce sparsity regularizer when we solve for parameter *δ*. In other words, we could append the *λ||β||*_1_ term to the end of Equation 5. However, since this method is less recognized in the field, we only focus on the former way to introduce sparsity through our experiment.

##### 2) Ridge Linear Mixed Model

Following how sparsity is introduced into LMM, another prominent regularizer, L_2_ norm regularizer, is naturally considered as a successor. Ridge Linear Mixed Model can be attained as following:

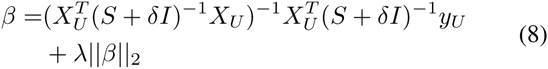

where *λ* is also a prior to control the strength of regularizer, as an extension from Ridge Regression [11].

Again, this regularizer can also be introduced directly to Equation 5 instead of Equation 6, but we adopt the above form for consistency with the sparsity case.

Different from the intuitive case of sparse Linear Mixed Model, Ridge regularizer may be expected to be less advantageous on variable selection performance. However, one strength is that L_2_ regularizer can help select the variables stably, while L_1_ regularizer usually sacrifice the stableness for sparsity [33].

##### 3) Other regularized linear mixed models

Many other regularizers can be introduced to Equation 6 or Equation 5 to aim for better variable selection performance or predictive performance.

For example, The L_0_ norm regularizer [15] has the ideal property to regularize effect sizes for a sparse variable selection. Unfortunately, its use forms a non-convex optimization problem, which is NP-hard [18], [6]. Some work aiming to combine the advantage of L_1_ norm and L_2_ norm. Elastic net is one such method [34], and Trace Lasso [9] improves Elastic net by offering the same modeling power with one fewer hyperparameter.

These regularizers can be straightforwardly applied to LMM. However, we do not cover them in this paper mostly because they are primarily the extensions of the classical L_1_ and L_2_ regularizers.

## III. MULTIPLE CONFOUNDING FACTORS CORRECTION

With the set-up of regularized linear mixed model, now we proceed to introduce different methods of handling multiple confounding factors. We will introduce three methods, namely Concatenation, Sequence, Interpolation. To be best of our knowledge, the Interpolation method has been explored, but not published^1^.

To better illustrate our idea, without loss of generalization, we assume the data sets come with two confounding factors, denoted as *Z*_1_ and *Z*_2_. These methods could be straightforwardly generalized to cases with finite number of confounding factors.

### A. Concatenation

The first method we present is simply the concatenation of confounding factors. Specifically, to achieve the same model as the standard one discussed previously with only one confounding factor, we simply concatenate two confounding factors into one, yielding

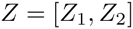

Therefore, kinship matrix *K* can be attained again as *ZZ*^*T*^.

This method may look trivial, but it effectively integrates different confounding factor categories into one universal category. For example, for a clinical study, if *Z*_1_ encodes that the subject is either *American*, *Asian*, or *African* and *Z*_2_ encodes the gender of a subject, then this concatenation method is indeed recasting *Z*_1_ and *Z*_2_ into *Z* that encodes that the subject is one of these six categories: *American male*, *American female*, *Asian male*, *Asian female*, *African male*, *African female*.

Therefore, we believe that concatenation is the simplest way of correct multiple confounding factors while maintains intuitiveness of integrating different categories into an universal one.

### B. Sequence

The second method we introduce is to correct multiple confounders following a sequential order.

For two confounding factors, *Z*_1_ and *Z*_2_, we have 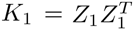 and 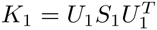,as well as 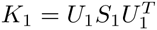 and 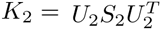

Then to correct multiple confounding factors in a sequential order, we successively transform the initial data (or current transformed data) into the next transformed space governed by the next confounding factor. In other words, we replace the term *U*^*T*^ *X* and *U*^*T*^*y* in Equation 5 and Equation 6 with 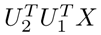 and 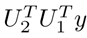, which could be trivially extended cases with infinite many confounding factors.

One fact to notice is that, *K* encodes the covariance structure of samples, therefore, for whatever information those confounding factors encode, *K* will be invariably of the same dimension as the number of samples available, leading to the fact all *U* matrix will of the same size, which validates the sequential equations above.

### C. Interpolation

The last method is an interpolation of available kinship matrices. Specifically, we first calculated 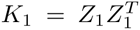 and 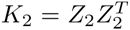 and we could derive *K* from a linear interpolation of *K*_1_ and *K*_2_.

For the simplest case, we have *K* = *K*_1_ + *K*_2_. This model makes the assumption that covariance structure is additive, therefore, a linearly interpolation of two covariance structure will result in a covariance structure that encodes both relationship information. This method will be helpful if two confounding factors are underlying related.

One simple question could be that the ideal interpolated result for *K* should be 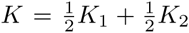 instead. However, since the first term learned by our model is *δ*, which is the scaling factor of covariance matrix, our model is consequently scale invariant of Kinship matrix. In other words, *K* = *K*_1_ + *K*_2_ will have the same statistical power with 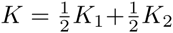.

## IV. EXPERIMENTS

In this section, we will investigate the performance of the regularized linear mixed models with different multiple confounding correction methods regarding variable selection and prediction on various data sets.

First, to fairly compare the methods, we generate a synthetic data set so that the golden standard result of variable selection is available as the evaluation metric. For this synthetic data set, the response variables are confounded with two different factors. Then, we create a semi-empirical synthetic data set where we use the real-world genome sequences as explanatory variables *X*. We generate effect sizes *β* and the response variable *y*, which is also confounded by two different sources of confounding factors. We evaluate the variable selection results with our generated *β*. Lastly, we evaluate the predictive performance with some real-world brain wave data relating to confusion mental state. With this data set, we compare our models on the F1 score of confusion prediction.

We compare the regularized linear mixed models with original un-regularized version when the parameters are learned through *MLE* or *REML* likelihood functions. The original linear mixed model, sparse linear mixed model and ridge linear mixed model are denoted as *linear*, *L*_1_ and *L*_2_ respectively.

We also compare the different methods we introduced in Section III. We investigate those three methods discussed together with the situation when we only correct one confounding factor. Therefore, there are five methods for the experiments and denoted as *Confounder 1*, *Confounder 2*, *Concatenation*, *Sequence*, *Interpolation* respectively.

### A. Synthetic Data Set

#### 1) Data Generation

Firstly, we generate explanatory variables, fixed effects and response variables that are corresponding to fixed effects. We generate a random matrix *X* with the size *n × p* and a random sparse vector *β* with the size *p ×* 1. Then we calculate vector *Y_f_* as *Xβ*, which is of the shape *n×*1. Therefore, we have explanatory variable *X*, fixed effect *β*. *β* serves as the golden standard of the variable selection task that our model is aimed to solve.

To simulate the situation where response variables are confounded by different confounding sources, we split *X* into two parts, the first half is denoted as *X*_1_ (of the size 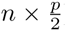) and the second half is denoted as *X*_2_ (of the size 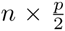). Then we cluster these *n* data points into groups based on the features *X*_1_ and *X*_2_ respectively by K-Means. For each group, we generate a corresponding response variable *Y_r_*_1_ and *Y_r_*_2_. The cluster information (i.e confounding factor identifiers) are *Z*_1_ and *Z*_2_.

Finally, we have the response variable as following:

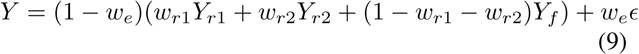

where ∈ is random noises that are sampled from a normal distribution *N*(0, 1). *w*_*r*1_ and *w*_*r*2_ are weights to control the strength of confounding factors. *w*_*e*_ is the weight to control the strength of random noises.

#### 2) Experiment Set-up

We compare three linear mixed models, under two different likelihood functions, *MLE* and *REML* respectively, and with three different multiple confounding factor correction methods as well as the situation when only one confounding factor is corrected.

Parameters of regularization for regularized linear mixed models are selected with cross-validation. We cross-validated the parameters *λ* to choose the model that identifies a fixed number (denoted as *M*) of causal variables [32]. The reason that we avoid traditional machine learning cross-validation mechanism that selects the parameter with highest predictive performance is that predictive performance is independent of the performance in variable selection.

We evaluate the performance with the precision-recall curve. As many different experiments are run, we show the comparison of area under precision-recall curve for simplicity. Table I reports the result under a various different choices of *M*.

#### 3) Results

Table I shows the area under the precision-recall curve for different models, different likelihood functions and various confounding factors correction methods under different reported *M* variables. The highest score of each row is indicated in italic and bold, while the second highest score is showed in bold only.

It is noticeable that for different confounding factor correction methods, *Sequence* method has a distinct advantage over others. Almost all the highest scores and second highest scores come from *Sequence* method.

To compare different regularized linear mixed models, we can see that each one of the three models has been reported as the highest score several times. However, sparse linear mixed models under sequence confounding correction method are consistently reported as the top two methods across all the numbers in each row. This correlates with the known fact that Lasso performs well in variable selection task due to its sparsity control.

We can also see that *MLE* and *REML* report identical scores. In fact, there are differences in almost all of these numbers if we look at the 7^th^ digit after the decimal point, but we cannot show these digits due to the limit of the page width. However, these differences are too insignificant, and we can safely draw the conclusion that the performances of *MLE* and *REML* are homogeneous.

**Table I:** AUC for precision-recall curve for different models, different algorithms and different confounding factor correction methods

| M | Confounder 1 |  |  | Confounder 2 |  |  | MLE<br>Concatenation |  |  | Sequence |  |  | Interpolation |  |  |
| --- | --- | --- | --- | --- | --- | --- | --- | --- | --- | --- | --- | --- | --- | --- | --- |
|  | Linear | L1 | L2 | Linear | L1 | L2 | Linear | L1 | L2 | Linear | L1 | L2 | Linear | L1 | L2 |
| 10 | **0.1034** | 0.0477 | 0.1033 | 0.1033 | 0.0477 | 0.1033 | 0.1033 | 0.0477 | 0.1033 | 0.1032 | **0.1588** | 0.0477 | 0.1033 | 0.1032 | 0.1032 |
| 50 | 0.1500 | 0.1394 | 0.1397 | 0.1499 | 0.1394 | 0.1397 | 0.1499 | 0.1496 | 0.1396 | 0.1398 | **0.1504** | **0.1598** | 0.1500 | 0.1293 | 0.1498 |
| 100 | 0.1292 | 0.1391 | 0.1340 | 0.1391 | 0.1391 | 0.1340 | 0.1290 | 0.1391 | 0.1289 | **0.1646** | **0.1449** | 0.1340 | 0.1241 | 0.1440 | 0.1290 |
| 150 | 0.1392 | 0.1324 | 0.1390 | 0.1392 | 0.1324 | 0.1356 | 0.1358 | 0.1291 | 0.1323 | **0.1500** | **0.1501** | 0.1490 | 0.1357 | 0.1292 | 0.1389 |
| 200 | 0.1319 | 0.1317 | 0.1318 | 0.1393 | 0.1293 | 0.1317 | 0.1417 | 0.1317 | 0.1317 | 0.1380 | **0.1431** | **0.1468** | 0.1318 | 0.1318 | 0.1317 |

| M | Confounder 1 |  |  | Confounder 2 |  |  | REML<br>Concatenation |  |  | Sequence |  |  | Interpolation |  |  |
| --- | --- | --- | --- | --- | --- | --- | --- | --- | --- | --- | --- | --- | --- | --- | --- |
|  | Linear | L1 | L2 | Linear | L1 | L2 | Linear | L1 | L2 | Linear | L1 | L2 | Linear | L1 | L2 |
| 10 | **0.1034** | 0.0477 | 0.1033 | 0.1033 | 0.0477 | 0.1033 | 0.1033 | 0.0477 | 0.1033 | 0.1032 | **0.1588** | 0.0477 | 0.1033 | 0.1032 | 0.1032 |
| 50 | 0.1500 | 0.1394 | 0.1397 | 0.1499 | 0.1394 | 0.1397 | 0.1499 | 0.1496 | 0.1396 | 0.1398 | **0.1504** | **0.1598** | 0.1500 | 0.1293 | 0.1498 |
| 100 | 0.1292 | 0.1391 | 0.1340 | 0.1391 | 0.1391 | 0.1340 | 0.1290 | 0.1391 | 0.1289 | **0.1646** | **0.1449** | 0.1340 | 0.1241 | 0.1440 | 0.1290 |
| 150 | 0.1392 | 0.1324 | 0.1390 | 0.1392 | 0.1324 | 0.1356 | 0.1358 | 0.1291 | 0.1323 | **0.1500** | **0.1501** | 0.1490 | 0.1357 | 0.1292 | 0.1389 |
| 200 | 0.1319 | 0.1317 | 0.1318 | 0.1393 | 0.1293 | 0.1317 | 0.1417 | 0.1317 | 0.1317 | 0.1380 | **0.1431** | **0.1468** | 0.1318 | 0.1318 | 0.1317 |

### B. Semi-empirical Synthetic Data Set

Because of the proliferate usage of Linear Mixed Model in genome-wide association studies (GWAS), our second experiment is aimed to evaluate the performance in variable selection for GWAS. To simulate the real genome data set as well as maintain the golden standard to be known, we generate the causal SNPs based on real SNP sequence. This is the reason that our second experiment is named as Semi-empirical Synthetic Data Set.

#### 1) Data Sets

We use the real genome information from a well-studied plant Arabidopsis thaliana. The Arabidopsis thaliana data set we obtained is a collection of around 1300 plants, each with around 215k SNPs [1]. For these SNPs, we randomly select 1000 causal SNPs and randomly generate their effect sizes following a Normal distribution in the same way as in the previous section.

We generate the response variables *Y* following the same condition as introduced in previous section following Equation 9.

#### 2) Results

This experiment is set up identically as the previous one except that now we report 1000 discovered SNPs. However, variable selection task on genome sequences is significantly more challenging from variable selection task on randomly generated data set due to Linkage Disequilibrium (LD) [24]. To account for the fact that discovering a SNP that is in LD with the actual causal SNP is equivalent to discovering the actual causal SNP statistically, we regard a discovery of SNP that is 200K bases away from the actual causal SNP as a true positive discovery.

The results are showed in Fig. 1. As the figure shows, in terms of both precision-recall curve and the area under precision-recall curve, sparse linear mixed model under the *sequence* multiple confounding factor correction method performs the best across all the combinations.

To compare the performance of different multiple confounding factor correction methods, we can see that *sequence* methods perform consistently better than all the other methods under each model or each likelihood function. Interestingly, on the model aspect, under all the other confounding factor correction methods, sparse LMM, which is expected to be the best one in variable selection task, behave significantly worse than the other two models.

Similar as previous case, there is no distinguishable difference between method *MLE* and method *REML*. Again, if we look further than the 6^th^ digit after decimal point, we will observe differences between these two methods.

### C. Brain Wave Confusion Prediction

Our third experiment is focusing on the predictive performance of regularized linear mixed model. Before the experiment, we need set up the background of how prediction is performed with linear mixed models.

#### 1) Prediction in Linear Mixed Models

The prediction of linear mixed model needs to be carried out differently from general machine learning models, for the reason that the effect sizes *β* are learned in a setting when *X* is transformed into *U*^*T*^ *X*. However, for a classification task, to maintain the labels valid, *y* remains untransformed. Consequently, the effect sizes are learned as a projection from the space *U*^*T*^ *X* to *y*, instead of from *X* to *y* (for general machine learning) or from *U*^*T*^ *X* to *U*^*T*^ *y* (for linear mixed model in regression)

Therefore, for testing data set with explanatory variables *X*_*te*_, *β* is not directly applicable. It is also unfeasible to directly use *U*^*T*^ *X*_*te*_ for many reasons, the simplest one will be mismatch of dimensions.

To carry out valid prediction, we make another assumption that states the Kinship matrix we have from training data set and the kinship matrix we calculate from testing data set are from the same distribution. Therefore when we project the testing data set with testing kinship matrix, it is still applicable to learned *β*.

In other words, we assume *Z*_*te*_ is available, and we have:

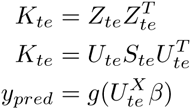

where *g*(*•*) stands for the function of classification (As an example, sigmoid function for Logistic Regression). Therefore, the predicted labels is attained from *y*_*pred*_

**Fig. 1:**
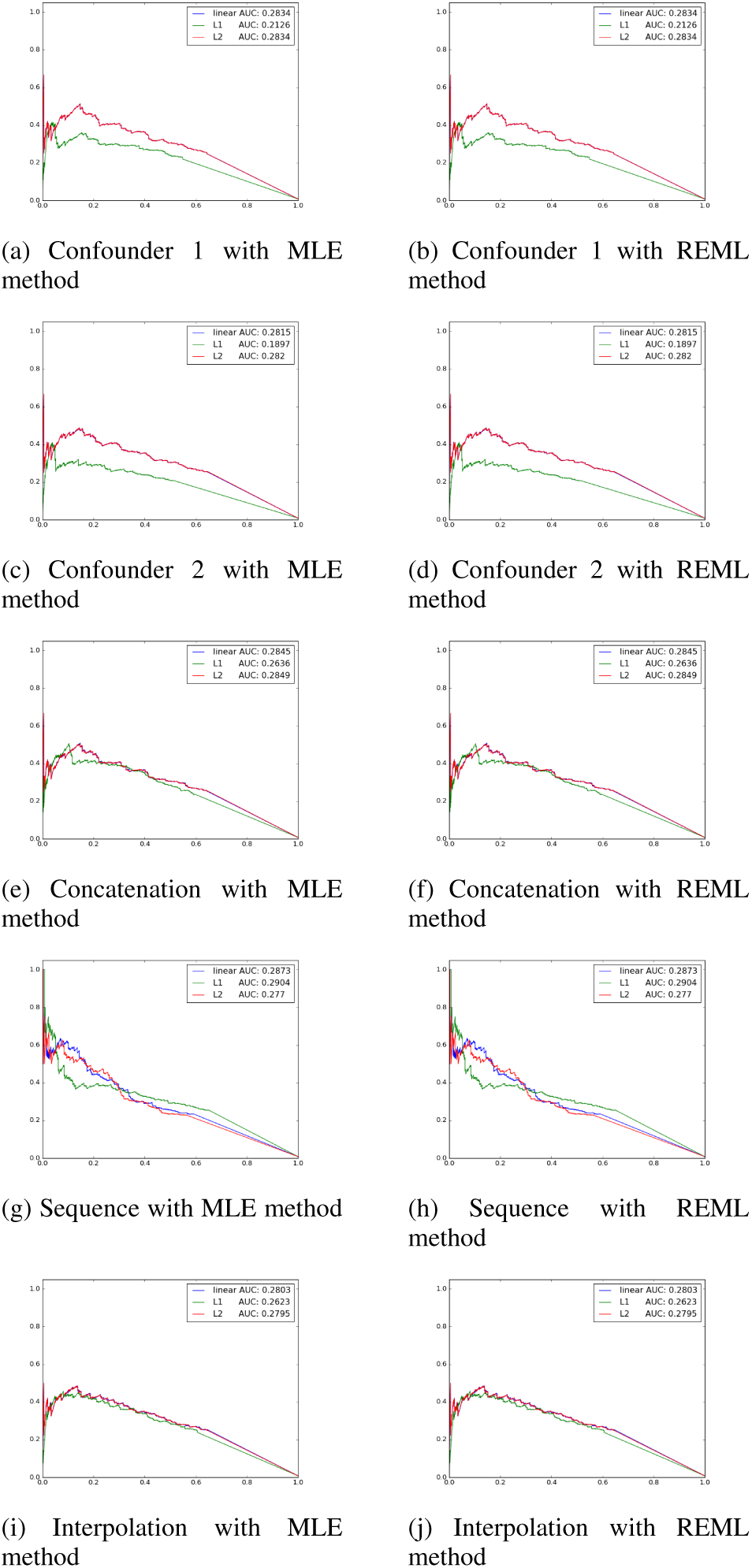
Precision-recall curve and AUC to compare the performance for each linear mixed model (in the same figure), for each learning algorithm (in the same row) and for each multiple confounding factors corrections method (in the same column)

#### 2) Data Set

We use the brain wave data set^2^ collected by EEG sensors when students are learning online courses [29]. The data is labeled as whether the students feel confused or not.

Data is collected from 10 college students while they watched MOOC video clips. Videos are extracted from online education resources that are anticipated not to be confusing for college students, as well as videos that are expected to confuse a typical college student if a student is not familiar with the video topics. Data is collected via a portable EEG mindset with a single sensor on the forehead.

We have 10 college students, each watching 10 videos and after they watch the video, they label that whether they feel confused or not themselves.

In our experiment, to make the task more challenging, we use the each sampled signal as one data point, resulting in more than 12k data points. Each data point shares the same label as the subject labels for that video. There are two confounding factors will highly affect the labels: i.e. subject IDs and video IDs.

To make the problem even more challenging, we use the first five subjects’ data as training data and the rest five subjects’ data to test. Therefore, the labels are heavily confounded by subject IDs since brain waves may be dramatically different between subjects and there is no data in the testing case of the subjects that appear in training case.

#### 3) Results

We compare the performance of regularized linear mixed models on methods handling multiple confounding factors on F1-score of confusion prediction. The results are showed in Fig 2.

As Fig 2 shows, both MLE and REML perform similarly in the learning process. Sparse Linear Mixed Model performs much better than the other two models. We conjecture that this is because of the overfit proof properties introduced by L_1_ regularizer.

For multiple confounding factors correction methods, we can see that *interpolation* works the best across these three methods. However, none of these methods work better than the simple case where we only correct confounding factor 1, which is the subject ID. We conjecture several reasons for this phenomenon: 1) the way we split training data and testing data makes the labels heavily confounded by subject IDs, but much less confounded by video IDs. 2) The labels we use are annotated by the subjects indicating whether they are confused or not. The labeling mechanism is naturally confounded by subjects IDs, but not video IDs. For these two reasons, we believe that it is reasonable to observe that the results behave the best when only the subject ID confounder is corrected, but not both.

## V. DISCUSSION

Through these three experiments, we have compared the performance of variable selection task and binary classification task for different linear mixed models, different likelihood functions and different multiple confounding factor correction methods. Several conclusions can be drawn when comparing these three experiments together.

**Fig. 2:**
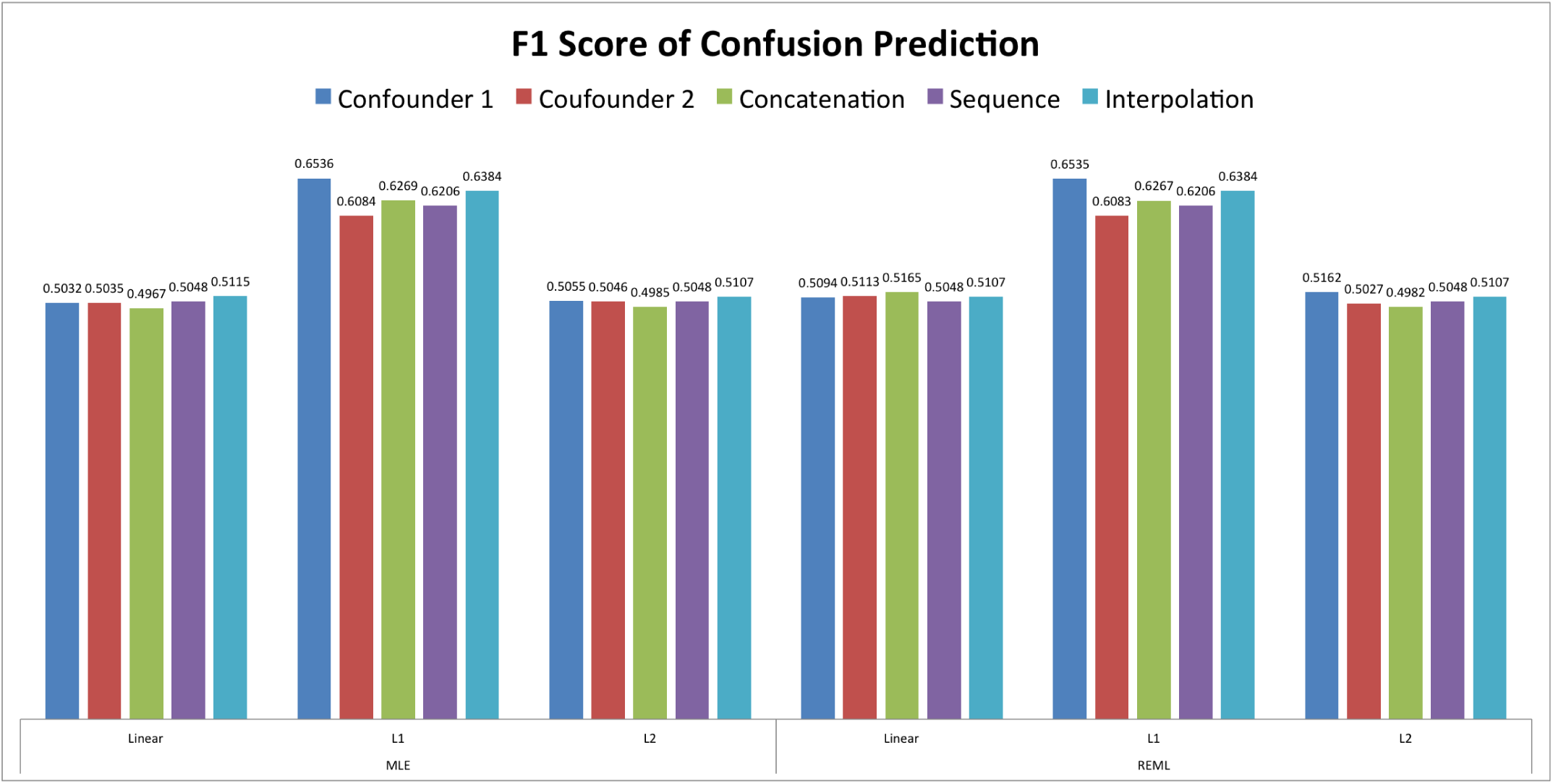
F1 score of predicted result of confusion

Sparse Linear Mixed Model under *sequence* confounding correction method consistently performs as the best (or at least second best) in variable selection tasks across these experiments. On one hand, while it is a known fact that L_1_ regularized models are good at variable selection tasks, it is interesting to see that Sparse Linear Mixed Model does not dominate other two models in other confounding correction methods. On the other hand, all these linear mixed models perform better with *sequence* method than corresponding models with other methods. We conjecture that, *sequence* method could correct those multiple confounding factors in a more ideal way, that it offers an excellent background for L_1_ regularized models to perform.

There is no observable difference between *MLE* and *REML*. Despite the fact that there are statistical differences, we do not observe any empirical differences when two methods are applied.

Selection of confounding factors correction models heavily relies on the semantics of confounding factors. In our third example, when we know the data is heavily confounded with the first confounders, and much less confounded with the second one, all these three multiple confounding factor correction methods perform inferior to the method when only the first factor is corrected.

Sparse Linear Mixed Models also dominate the performance with respective to prediction. This is probably related to the overfitting proof property introduced by L_1_ regularizers.

## VI. CONCLUSION

In this paper, we discussed regularized linear mixed models with L_1_ and L_2_ regularizers, reviewed two methods that used in parameter learning for linear mixed models, namely *MLE* and *REML* and then introduced three methods to correct multiple confounding factors together.

We evaluated the performance for two tasks that are frequently met in biomedical research, namely variable selection task and prediction task. We first evaluated the methods on synthetic data. Then we evaluated the methods on the semi-empirical data set generated from genome information to simulate GWAS, which was followed by evaluation of prediction performance on brain wave data set that is related to the mental state of confusion.

Our experiments have shown that Sparse Linear Mixed Models under *sequence* method of multiple confounding factors correction performed the best consistently compared across all the other combinations. *MLE* and *REML* methods did not distinguish from each other. We also made some suggestions on selection of multiple confounding factor correction methods.

Looking forward, we would like to explore more on the theoretical aspect of those multiple confounding factor correction methods and build the selection guidance of these methods in the theoretical aspect in addition to the empirical aspect discussed in this paper.

## ACKNOWLEDGMENT

The authors would like to thank Pittsburgh Super Computing Center for the computation resources. We would also like to thank Professor Daniel Weeks from University of Pittsburgh and Professor Howard Seltman from Carnegie Mellon University for suggestions on presentation of this paper.

## Footnotes

1 https://github.com/MicrosoftGenomics/FaST-LMM/tree/master/fastlmm

2 Publicly available at: https://www.kaggle.com/wanghaohan/eeg-brainwave-for-confusion

